# Dual Adversarial Deconfounding Autoencoder for joint batch-effects removal from multi-center and multi-scanner radiomics data

**DOI:** 10.1101/2023.01.16.524181

**Authors:** Lara Cavinato, Michela Carlotta Massi, Martina Sollini, Margarita Kirienko, Francesca Ieva

## Abstract

Medical imaging represents the primary tool for investigating and monitoring several diseases, including cancer. The advances in quantitative image analysis have developed towards the extraction of biomarkers able to support clinical decisions. To produce robust results, multi-center studies are often set up. However, the imaging information must be denoised from confounding factors – known as batch-effect – like scanner-specific and center-specific influences. Moreover, in non-solid cancers, like lymphomas, effective biomarkers require an imaging-based representation of the disease that accounts for its multi-site spreading over the patient’s body. In this work, we address the dual-factor deconfusion problem and we propose a deconfusion algorithm to harmonize the imaging information of patients affected by Hodgkin Lymphoma in a multi-center setting. We show that the proposed model successfully denoises data from domain-specific variability while it coherently preserves the spatial relationship between imaging descriptions of peer lesions, which is a strong prognostic biomarker for tumor heterogeneity assessment. This harmonization step allows to significantly improve the performance in prognostic models, enabling building exhaustive patient representations and delivering more accurate analyses. This work lays the groundwork for performing large-scale and reproducible analyses on multi-center data that are urgently needed to convey the translation of imaging-based biomarkers into the clinical practice as effective prognostic tools. The code is available on GitHub at this [link](https://github.com/LaraCavinato/Dual-ADAE)

## 1. Introduction

Hodgkin Lymphoma (HL) is a type of cancer that affects the lymphatic system, where lymphocytes proliferate uncontrollably in multiple lymph nodes and eventually in extranodal sites (e.g. spleen, bone, etc.). It is acknowledged as a curable disease thanks to its high rate of response to chemotherapy, often combined with radiotherapy. Still, a considerable percentage of patients do not respond to first-line treatments and the latest research has been devoting its efforts to discovering alternative and more efficient therapies, such as immunotherapy. Immunotherapy has indeed been approved for relapsing cases and has since represented a huge stride for patients, who are on average very young (Mohty et al., 2021).

As the number of available therapies increases, treatment planning becomes more and more crucial, and personalized medicine is catching on in every aspect of medical practice to devise the optimal treatment for each patient. Nevertheless, such a tailored approach requires quantitative and informative data to input into powerful and transferrable models on which to rely decisions. On purpose, Positron Emission Tomography/Computed Tomography (PET/CT) radiomic analysis has been shown to be an insightful, non-invasive tool for histological prediction, prognostic assessment, and bone marrow involvement definition in lymphoma (Rizzo et al., 2021). In brief, the radiomics framework entails the extraction of a high-dimensional vector description of the spatial gray levels’ distribution of an image, the so-called radiomic features (Afshar et al., 2019, Scapicchio et al., 2021). Each of such features thus describes a statistical property of the image heterogeneity at different scales, which can inform several downstream analyses and modeling efforts.

As HL is a rare disease, studies performed at a single institution usually do not account for sufficient information to build powerful enough models and derive general knowledge. Therefore, oftentimes multi-center cohorts need to be set up and large-scale studies have to be conducted, collecting data coming from different sources (Parmar et al., 2018). This raises a relevant issue, as radiomics features are known to be highly influenced by the image acquisition settings and the reconstruction parameters, jeopardizing the transferability and scalability of the results (Berenguer et al., 2018, Pavic et al., 2018). Typical exogenous confounding factors include both scanner characteristics and protocols and more general center-specific variabilities. These two factors must therefore be accounted for together when performing any type of analysis on multi-center data.

Moreover, the latest trend in radiomics is developing towards the extraction of more and more features, including first-order statistics, second- and higher-order statistics, and wavelet/frequency-derived indices. As the number of features rises, their pairwise correlation increases accordingly, and it becomes harder and harder to build effective models and disentangle the true signals of interest from technical artifacts, noise, and uninteresting biological variables. Here comes the need to properly reduce the dimensionality of radiomics vectors, transforming the features into low-dimensional vectors that keep the true informative signals while discarding domain-specific confounders. That is, both scanner and center-related variability must be removed from radiomic features leaving the predictive tumor-specific information intact.

While the above holds for many multi-center radiomics studies of (rare) diseases, when analyzing a hematological (like HL) or metastatic cancer, an additional level of complexity is added to the task of deconfounding and reducing radiomics features. In fact, different lesions can be found throughout the body of the patients. Despite the current approach for imaging-based quantitative assessment of most cancers, including HL, relies on the inspection of the bigger or hotter lesion, Sollini et al. (Sollini et al., 2020) have demonstrated how lesions are radiologically heterogeneous within patients in terms of radiomics description and how a prognostic classifier performs better when all tracer-avid lesions are considered. These findings align with the latest discoveries in the biological underpinnings of lymphomas. Some studies on solid cancers have previously described how both proximal and distant lesions deriving from the same primary tumors exhibit divergent patterns of both morphological and genetic heterogeneity (Campbell et al., 2008). Similarly, Tabanelli et al. (Tabanelli et al., 2020) reported the same evolutionary crossroad between morphological heterogeneity and intra-clonal evolution in a case of high-grade B-cell lymphoma. Thus, morphological heterogeneity behaves as a surrogate of genetic heterogeneity, responsible for treatment inefficacies. It follows that all lesions’ morphology must be taken into account, to exhaustively represent the disease in the prediction of cancer progression, therapy efficacy, and disease-free survival outcomes (Sangaletti et al., 2020, Lavin and Tan, 2022). This implies that any postprocessing (i.e. dimensionality reduction and/or deconfusion process) aimed at preparing radiomics features for patients’ representations needs to keep the inter-lesion relationships within patients consistent, as here is where information of tumor morphological heterogeneity lies (Sollini et al., 2020, Cavinato et al., 2022).

In light of the above, a robust post-image-acquisition method aimed to harmonize multi-lesion radiomics data from multi-center studies requires (i) to properly remove both scanner and center confounding effects, (ii) to treat features’ collinearity and allow for simpler statistical modeling via proper dimensionality reduction, and (iii) to keep intra-patient heterogeneity consistent throughout the transformation. All this should be achieved while retaining all truly informative signals in the data, so as not to affect – and possibly improve - any potential down-stream analysis.

Different strategies have been proposed in recent literature to minimize the batch-effects of radiomics variability, ranging from imaging-based to feature-based approaches (Da-Ano et al., 2020, Ligero et al., 2021, Mali et al., 2021). Most of them aim to perform batch-specific standardization of images to disentangle the true signal from environment-related noise. Among these, the ComBat method was shown to be superior to other techniques, attracting attention in the radiomics field (Johnson et al., 2007, Chen et al., 2011, Da-Ano et al., 2020). ComBat is a statistical harmonization method that can correct variation by using empirical Bayes to estimate location and scale parameters. Starting from its first conception, ComBat was improved over time by different independent researchers. One for all, Adamer et al. proposed a regularized solution of ComBat, namely ReComBat, computationally more efficient to facilitate the large-scale harmonization of data (Adamer et al., 2022). However, it must be noted that ComBat and most of its derivative algorithms were developed in the computational biology domain, where usually only one main confounder (i.e. sequencing batch effect) needs to be removed. Indeed, to remove multiple confounders, they must be applied repeatedly, one factor at a time. As the context of radiomics studies oftentimes implies multiple confounders, Nested ComBat (Horng et al., 2022b) and its improved evolution from the same authors, OPNested Combat (Horng et al., 2022a), were recently proposed specifically to tackle multi-factor deconfusion. The latter applies ComBat iteratively on confounder-associated subsets of features, identifying the optimal order of factors to correct for. Notably, irrespective of the number of confounders removed from the data, ComBat-based methods rely on the hypothesis of normality of the features’ errors, which might be unrealistic for radiomics data (Horng et al., 2022a). Moreover, none of the above methods perform dimensionality reduction and are thus typically followed by Principal Component Analysis (PCA) before the analysis. Additionally, to the best of our knowledge, none of them has neither explicitly addressed the problem of preserving inter-lesion relationships within patients, nor has been evaluated in their capability to improve prediction by exploiting heterogeneity information after deconfusion.

In this work, we propose a multi-factor deconfusion algorithm better suitable for the downstream analysis of multi-lesion/metastatic patients in multi-center studies. The algorithm builds on the work of Dincer et al. (Dincer et al., 2020), which, in the context of gene expression analysis, proposes an Adversarial Decon-founding AutoEncoder (AD-AE) model that requires no assumption on features’ distribution and jointly performs dimensionality reduction and cleaning of the embeddings, enhancing the signal-to-noise ratio. Here, we exploit the rationale of this model for the context of multi-center PET/CT radiomics analysis, developing a dual factor AD-AE (in the following, Dual AD-AE) model for the simultaneous removal of both center and scanner confounding effects. We evaluate the proposed model in terms of (1) its deconfusion power, (2) its ability to keep invariance of intra-lesion relationship with respect to original data - despite dimensionality reduction - (3) and its prognostic power. In experiments (1) and (3) we compare the results of Dual AD-AE to those of state-of-the-art ComBat-based approaches. In experiment (2), we propose a statistical test to access the consistency of the data transformation. We evaluate our proposed models on a multi-center dataset of HL patients in order to predict response to first-line chemotherapy, demonstrating that Dual AD-AE enables building exhaustive patient representations and delivering more accurate analyses, especially when trying to exploit the predictive power of intra-tumor heterogeneity.

## 2. Methods

In this section, we describe the models and the experimental procedures used in this study. Specifically, we include a description of the study population and the sampling method, as well as the instruments or tools used to collect the data. Additionally, we provide a mathematical overview of the proposed models along with state-of-the-art approaches. We then further detail the design of the experiments, stating the hypotheses and the research questions that the study aims to address.

### 2.1. Data collection

Two centers were involved in the study; inclusion criteria were age ≥ 16 years old, newly diagnosed stage I-IV HL, baseline [^18^F]FDG-PET/CT availability, and exclusion criteria were missing clinical/imaging/follow-up data; 128 HL patients were recruited and treated at IRCCS Humanitas Research Hospital (Institution 1), 78 at Fondazione IRCCS Istituto Nazionale dei Tumori (Institution 2). Personal information and clinical data were annotated for each patient in both hospitals and [^18^F]FDG PET/CT imaging was inspected by experienced nuclear medicine physicians. Descriptive statistics of patients are available in Table A.3 and Table A.4 (Appendix A) for Institution 1 and Table A.5 and Table A.6 (Appendix A) for Institution 2. All [^18^F]FDG-avid lesions bigger than 64 voxels were segmented in each patient and radiomic features were extracted from each lesion using LIFEx software (Nioche et al. 2018, www.lifexsoft.org). A total of 1340 and 794 lesions were collected at Institution 1 and Institution 2, respectively. Information about scanners’ specification and acquisition settings is summarized in Table B.7 and Table B.8 (Appendix B), while Imaging Biomarker Standardization Initiative (ISBI)-compliant standardization and data harmonization have been published elsewhere (Sollini et al., 2020). The study was approved by the local ethics committees at Institution 1 (n. 2595 on Jun16, 2020) and Institution 2 (code INT 212/20 on Sep28, 2020); given the observational retrospective design of the study, the signature of a specific informed consent was waived.

### 2.2. Dual Adversarial Deconfounding Autoencoders

We propose Dual Adversarial Deconfounding AutoEncoders (Dual AD-AE) to jointly tackle the denoising from both center- and scanner-related information. The architecture of the Dual AD-AE is described in Figure 1. The network consists of two parts: one autoencoder and an adversary branch. The autoencoder takes as input the radiomic vector associated with a lesion and performs the dimensionality reduction. It is made of one input layer (number of input nodes: [1 × 45]) two hidden layers (number of first hidden layer nodes: [1 × 32], number of second hidden layer nodes: [1 × 16]), and one output layer (number of output nodes: [1 × 45]). The autoencoder represents the backbone of the model and, from its deepest layer, two adversary networks branch out for center and scanner predictions. Both adversary networks are made of two hidden layers (dimensions of the first hidden layer and the second hidden layer are [1 × 50] and [1 × 50] respectively) and one output layer ([1 × 2] for center prediction and [1 × 5] for the branch predicting the scanners).

**Figure 1:**
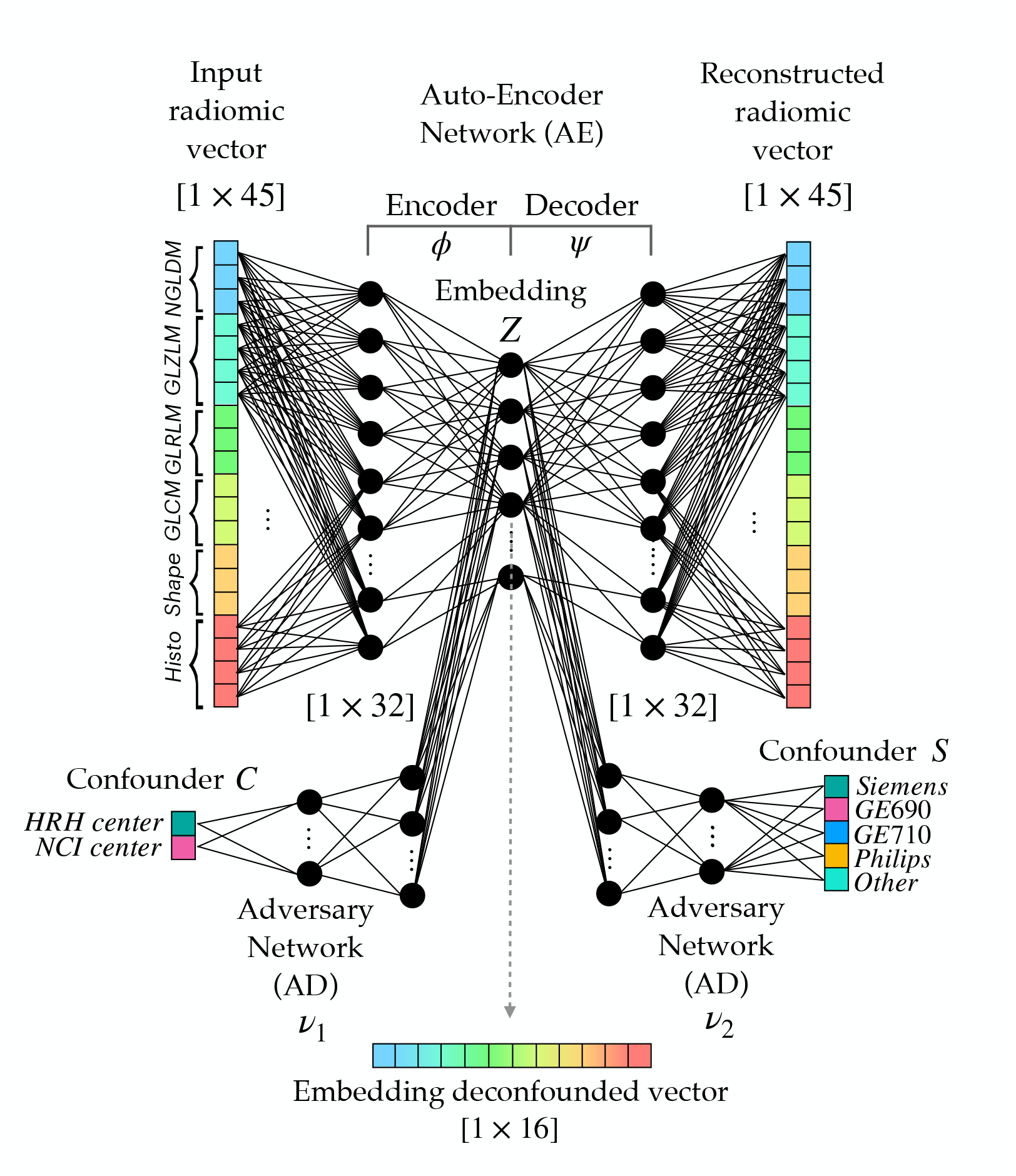
Architecture of the Dual Adversarial Deconfounding Autoencoder (AD-AE) model: the model is made of three parts: an autoencoder (encoder: *ϕ*, decoder: *ψ*), an adversary branch network predicting the center confounder (*v*_1_) and a parallel adversary branch network predicting the scanner confounder (*v*_2_). The network is trained by optimizing the input reconstruction task (autoencoder loss) and the deconfusion task (adversary losses) as in Equation 1. The adversaries unlearn to predict the confounding factors, i.e. the center and the scanner.

The loss is then made of three terms, where the reconstruction error, the accuracy of the center classification, and the accuracy of the scanner classification sum up as in Equation 1:

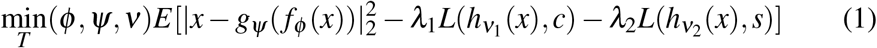

where *v*_1_ is the center adversary branch, *v*_2_ is the scanner adversary branch, *λ*_1_ and *λ*_2_ are weighting parameters and *c* and *s* are the true labels for center and scanner respectively. Of note, weighting parameters can be tuned to tailor the importance of the tasks to be optimized. For instance, one could prioritize one confounding factor rather than the other, having *a priori* information about the latent variability of the specific case study data.

In our setting, hyperparameters were tuned according to grid search. The number of layers, the number of nodes, and weighting parameters were optimized based on the reconstruction error. The number of epochs was optimized according to early stopping strategy (Yao et al., 2007), i.e., iterations were stopped when no relevant improvements of the validation loss were recorded. The batch size was set to 128 and *λ*_1_ = *λ*_2_ = 1.

### 2.3. Benchmark state-of-the-art Methods

Among the methods proposed in the literature for imaging harmonization, ComBat has been repeatedly elected as the best approach such that different implementations and further improvements have been proposed in the last years.

ComBat was originally proposed by Johnson and Rabinovic (Johnson et al., 2007) for removing the batch-effect seen in genetics microarray analysis. The harmonization method consists of standardizing each batch according to its mean and variance. Specifically, the correction takes place at a specific location and scale (*L/S*), wherein the batch-related error is supposed to be present. *L/S* model states that the value *Y* for feature *f* from sample *j* in batch *i* follows the following formulation:

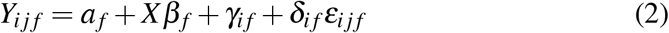

where *a_f_* is the feature value, behaving as intercept; *X* is the design matrix and *β_f_* is the features coefficients such that *Xβ_f_* is the observed variability; *γ_if_* and *δ_if_* are the additive and multiplicative batch effects respectively and *ε_ijf_* the standard error. Accordingly, *γ_if_* and *δ_if_* can be estimated (either in parametric and non-parametric ways) from data, and *Y_ijf_* can be corrected as:

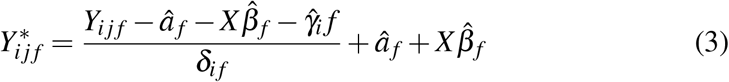

One of the main advantages of ComBat is being effective even with small batch sizes. Being 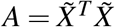 positive-definite, the optimization problem is strictly convex. However, when *A* happens to be singular the regression estimation does not exist and, if the system is underdetermined, ComBat does not guarantee to bring out a unique solution. For this reason, Adamer et al (Adamer et al., 2022) proposed a regularized solution of ComBat (ReComBat) computationally more efficient to facilitate the large-scale harmonization of data.

Applying ComBat or ReComBat in cascade to capture and remove the linear variability from more than one confounding factor may cause instabilities depending on the specific order of the harmonization steps. Very recently, Horng et al (Horng et al., 2022a,b) proposed an optimized procedure for sequentially harmonizing data from multiple batch effects, namely OPNested ComBat. In our experimental setting, besides ComBat and ReComBat, we included OPNested as a benchmark model, to be tested in both deconfusion and predictive powers.

### 2.4. Evaluating Point-Cloud shape consistency

As stated in Section 1, a solid harmonization method for multi-lesion patient representation needs to keep invariance of intra-lesion relationship with respect to original data, even after the dimensionality reduction step. In multi-lesion and/or metastatic tumor settings such as HL, patients can be modeled as clouds of points (Cavinato et al., 2022), where each point is defined by the radiomic vector – whether original, reduced, or deconfounded – of a lesion, and the shape of the cloud determines intra-patient tumor heterogeneity as the pairwise relationship between lesions (Gil et al., 2021). To test the point-cloud shape consistency across transformation, we here define a novel approach: the Point Cloud Semantic Drift (PCSD).

Before defining PCSD, let us introduce some necessary notation. Let *M_i_*(1)…*M_i_*(*K*) be the scores associated with the ordered list *L_i_*, where *M_i_*(1) is the best score, *M_i_*(2) is the second best, and so on. The best score can be the largest or the smallest depending on the context. Let 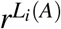 be the rank of *A* in the list *L_i_* if element *A* is within the top *k* elements, and be it equal to *k* + 1 otherwise; *r^δ^*(*A*) is defined likewise for a different list *δ*. The Spearman’s footrule distance between *L_i_* and any ordered list *δ* can be defined as:

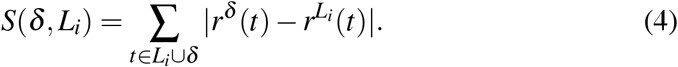

Equation 4 is the sum of the absolute differences between the ranks of all the unique elements of the union of the two ordered lists. The smaller the value of the metric, the more similar the lists. To compute the Point Semantic Drift (PSD) for an arbitrary point *t*, we exploit a weighted version of *S*. We estimate the PSD as the weighted change in neighbor rankings, according to Equation 5.

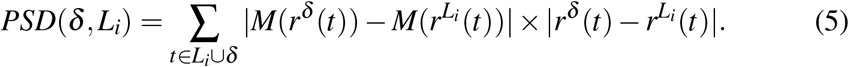

PSD is the sum of penalties for moving an arbitrary element (data point) *t* of the list *L_i_* from a position *r^δ^*(*t*) to another position 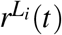 within the list (second term of the product) adjusted by the difference in scores between the two positions (first term).

*M*(*r^δ^*(*t*)) and 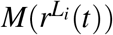 are the normalized distances between *t* and all other points in the cloud, after and prior to any transformation, respectively. This weighting scheme penalizes more the changes in the positions of very distant points, than the neighboring shifts of observations lying close in the original cloud. That is, higher weights are assigned to swaps between close-by and far-distant points, compared to changes among close neighbors.

Once computed the PSD for each point in the cloud *C*, the Point Cloud Semantic Drift is estimated as the average PSD_*k*_ of the *K* points in *C*:

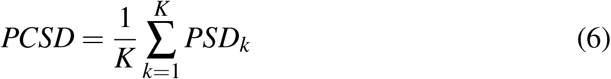

where *K* is the number of lesions in the patient under consideration.

In our setting, *L_i_* corresponds to the set of lesions of patient *i* as described by the raw radiomic features (original set); *δ* corresponds to the set of lesions described by the transformed features after deconfusion (e.g. Dual AD-AE mode). PCSD thus accesses and quantifies the invariance of each cloud (patient) to the data transformation process.

Given that PCSD can take on values ranging from 0 to infinity, we need to establish a suitable test to assess the significance of the results obtained from our deconfounded point clouds. To accomplish this, we can build a null distribution of PCSD values (PCSD*null*) which serves as an upper bound for the drift. That is, it represents the change in the cloud’s shape that would occur if an arbitrary embedding function was employed, completely disregarding the initial data structure. Operationally, we randomly transform the original cloud by adding a random Gaussian noise with mean *μ* = 0 and variance *σ* = 3 to a different subset of the lesions’ vectors. We do this iteratively for 100 times, each time computing the PCSD each time. Upon these values we build the Empirical Cumulative Distribution Function. If the true PCSD value obtained from our deconfounded embeddings fall within the limits of the left tail of this empirical null distribution, significant evidence is obtained on the ability of our algorithm to maintain the original cloud structure. The empirical p-value is computed from the Empirical Cumulative Distribution Function by computing the ratio between the number of trials where the PCSD is lower than the current and the total number of trials (100).

### 2.5. Experimental outline

To test the robustness of any post-image-acquisition method aimed at harmonizing multi-lesion radiomics data from multi-center studies, one needs to consider several aspects. We performed a series of experiments on different harmonization strategies (in the following, *modalities*), including our proposed Dual AD-AE model and different ComBat-based methods.

We recall that the center confounding factor relates to the hospital’s imaging facility, the clinical guidelines, and the personnel who segments and carries out the acquisition. On the other hand, the scanner confounding factor supplies information on the scanners’ specifications and reconstruction parameters. The scanner variable is intrinsically subordinated to the center variable, as usually different scanners are found in different centers. That is, the two sources entail some extent of nesting nature and may then partially overlap in their confounding information.

For the sake of comparison with state-of-the-art approaches, we tested three major ComBat implementations, namely Combat (Johnson et al., 2007), ReCom-Bat (Adamer et al., 2022) and OPNested Combat (Horng et al., 2022a), comparing the results to quantify the improvements of our solution. We employed different pipelines to test their performance from different perspectives. ComBat was used for deconfounding the imaging data from the center and scanner information. The two ComBat models were applied in cascade to the data: (1) one label was used as a batch effect to be removed and (2) the obtained denoised vector was further deconfounded by the effect of the other label. We followed two different orders, namely ComBat-center-scanner and ComBat-scanner-center. The very same procedure was investigated by employing ReComBat implementation. Two different pipelines were thus derived, namely ReComBat-center-scanner and ReComBat-scanner-center. OPNested Combat was instead applied once on center and scanner effects at the same time, as it was specifically developed for multi-factor effect removal.

On these models, we performed three different quantitative experiments. We tested the deconfusion power of the different modalities, comparing the proposed method to the state-of-the-art models (Experiment 1). Furthermore, considering that the Dual AD-AE encompasses dimensionality reduction as part of the deconfusion process, leading to a potentially detrimental transformation of intra-lesion relationships, we implemented our PCSD test to assess this impact quantitatively (Experiment 2). Finally, we tested and compared all modalities on their ability to keep predictive information intact. We transformed the deconfounded features of each modality into different all-lesions patients’ representations, to be fed into prognostic models (Experiment 3). Details are provided in the following.

#### 2.5.1. Experiment 1

To evaluate the strength of the confounders’ effect, one can verify the predictability of the confounding variables (i.e. the center and the scanner) starting from the data. A high prediction performance denotes the presence of a strong confounder-related signal. Therefore, to quantify the effect of the deconfusion process, we employed a cross-validated Logistic Regression model, with 100 trials and replacement. Testing accuracy was annotated in each trial to compute the mean trend and the standard deviation of the performance of each modality. Additionally, to compare the performance of the models, given the normality of the data, we used a two-sided parametric t-test for paired samples and evaluated the improvements of the different harmonization strategies compared to the pure radiomics description.

#### 2.5.2. Experiment 2

To ensure that the information on the clouds’ shape has been preserved, Dual AD-AE embeddings must keep invariance concerning the relative positions of lesions, despite the reduced dimensionality of the resulting vectors. Patient-wise lesions’ rankings and pairwise lesions’ distances should hold after deconfusion, under the hypothesis that they are not independently impacted by exogenous noise. Given this assumption, to test for the cloud shape-invariance of the feature transformations, we implemented the Point Cloud Semantic Drift (PCSD) quantitative method. As explained in Section 2.4, PCSD quantifies the extent of the change in peer lesions’ distance rank order within a patient. Furthermore, to define a quantitative test of hypothesis for PCSD, we estimated an empirical null distribution of the PCSD values when point clouds are transformed at random, inducing random neighbor swaps by injecting repeatedly Gaussian noise in subsets of the embeddings. The PCSD was computed for each patient and a population test for testing the transformation consistency was carried out in the context of the Dual AD-AE. The empirical p-values of the Dual ADAE transformation were then obtained from the Empirical Cumulative Distribution Function (ECDF) of this null PCSD distribution.

#### 2.5.3. Experiment 3

Despite its unsupervised nature, the proposed approach aims to enable the design of exhaustive patient representations to deliver accurate analyses for treatment planning on multi-center datasets. Here we provide an example of downstream analysis where to quantify the improvement in predicting the first-line chemotherapy outcome of patients affected by HL after correcting for confounding factors. Specifically, the predictive power of the imaging features has been evaluated with Cox proportional hazard survival models in a cross-validation fashion.

Three patient representation strategies were implemented to summarize the multi-lesion information in a single vector object to be properly fed into the models. First, the centroid of each patient’ point cloud was computed as the mean profile of peer lesions belonging to them (“centroid representation”). Then, as a second patient representation, only the distribution of the lesions over the statistical space was described and used as model input. For each patient, we computed the pairwise distances between all lesions in the patient and we calculated the mean and the standard deviation as indexes for lesions’ variability. Moreover, we took the distances between every lesion of the patient and their centroid and kept the average and the standard deviation of these distances to quantify the lesions’ spreading from their center. Thus, the four indexes were exploited as “point cloud description representation” to be fed into the survival model. Finally, the two abovementioned representations were merged in a “complete representation” of the patient encompassing both the mean disease profile of patients and the variability of their lesions. For each of the modalities under testing, the three representations were computed and fed into a Cox proportional hazard model (Lin and Wei, 1989) to predict the time-varying response to therapy. Of note, raw radiomic, ComBat- and ReComBat-based standardized radiomic features were reduced using PCA prior to being input to any model. To result in a dimensionality comparable to the embeddings, we kept the first sixteen principal components, accounting for at least 90% of the variability. Training and testing sets were repeatedly split multiple times (20 splits) and Concordance Index (CI, Harrell et al. 1982) scores were reported to assess the improvements that the harmonization step brings in terms of prognostic power. To do this, given the normality of the data, one-sided parametric t-tests for paired samples were employed to establish the optimal harmonization strategy. Specifically, the Dual AD-AE embeddings’ performance was compared to ComBat-center-scanner, ComBat-scanner-center, ReComBat-center-scanner, ReComBat-scanner-center, and OPNested ComBat.

## 3. Results

Three tests have been implemented to test for (1) deconfusion power, (2) transformation consistency, and (3) predictive power of the proposed algorithm compared to the current literature.

### 3.1. Experiment 1: Checking deconfusion power

Table 1 shows the results for the dual AD-AE, the two ComBat models, the two ReComBat models, and OPNested ComBat.

**Table 1:** Experiment 1 results: comparison between the performance of the Logistic Regression models for predicting the two confounding factors: the center and the scanners. The modalities that have been evaluated are raw radiomic data, Dual AD-AE embedding, ComBat-standardized data (both with center-scanner order and with scanner-center order), ReComBat-standardized data (both with center-scanner order and with scanner-center order) and OPNested-standardized data (with scanner-center order). The Logistics Regression models are fitted on each of these modalities, in a cross-validated fashion. Values are annotated as mean *pm* standard deviation. The models evaluate (1) the binary prediction of the center labels, and (2) the multi-class prediction of the scanner labels. The performances of the radiomics-based models are taken as reference, while the performances of the other modalities are analyzed in terms of decrease compared to the baseline models’ performance. Statistical tests have been performed and the models that are significantly different from radiomics are highlighted in bold.

|  |  | Accuracy CENTER | Accuracy SCANNER |
| --- | --- | --- | --- |
| **Baseline** | Radiomics | $0.8559 \pm 0.0117$ | $0.8617 \pm 0.0104$ |
| **Embedding** | Dual AD-AE | **$0.6251 \pm 0.0131$** | **$0.3308 \pm 0.0146$** |
| **ComBat** | ComBat-center-scanner | **$0.6220 \pm 0.0137$** | **$0.2968 \pm 0.0278$** |
|  | ComBat-scanner-center | **$0.6236 \pm 0.0134$** | **$0.3006 \pm 0.0307$** |
| **ReComBat** | ReComBat-center-scanner | **$0.6228 \pm 0.0150$** | **$0.3009 \pm 0.0362$** |
|  | ReComBat-scanner-center | **$0.6276 \pm 0.0124$** | **$0.2997 \pm 0.0359$** |
| **Opnested** | Opnested (scanner-center) | **$0.6239 \pm 0.0118$** | **$0.2967 \pm 0.0349$** |

While radiomics, as expected, scored very high in predicting both the center and the scanner, the embedding showed evidence of deconfusion, comparable to state-of-the-art benchmarks. Both the Dual AD-AE and all the Combat-based modalities aligned to the same performance, outperforming the non-deconfounded radiomic vectors. Indeed, values highlighted in bold in Table 1 correspond to non-significantly different, yet lower than radiomics, performances. All modalities were thus equally powerful at the deconfusion task. Of note, the OPNested algorithm selected scanner-center as the optimal order, thus the two models are expected to perform similarly. Additionally, the proposed Dual AD-AE model showed a smaller standard deviation of the accuracy in predicting scanner type, supporting the robustness of the model.

### 3.2. Experiment 2: Cloud-shape invariance test

Figure 2 shows the results of the proposed method. The population distribution of PDSC from Dual AD-AE transformation is displayed alongside the Empirical Distribution Function (EDF) of 100 random transformations. From the visual inspection of the plots, the model produced PCSD values skewed toward zero, suggesting the shape-invariance of the clouds. Moreover, the empirical p-value was equal to zero, thus we can further sustain that Dual AD-AE successfully kept cloud-shape invariance and that the change in inter-lesion distance, which occurred during deconfusion, was significantly far from being random.

**Figure 2:**
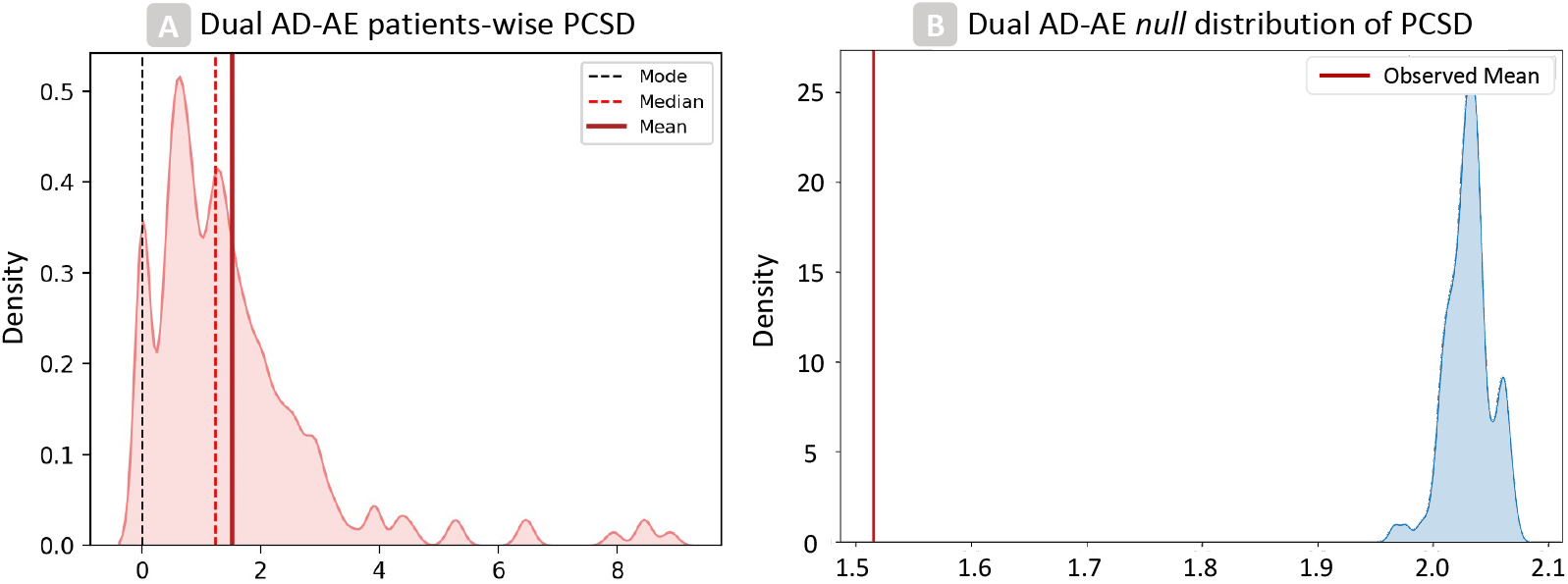
Results of patient-wise tests on PCSD for dual AD-AE embedding: (Panel A) The density plot displays the overall distribution of PCSDs in the population. (Panel B) The density plot shows the score of the dual AD-AE results over a bootstrap random distribution. Fiducial values are marked with vertical lines in the left plot and our model performances are displayed with a vertical red line in the right plot.

### 3.3. Experiment 3: Checking prognostic power

Performances on both the training and testing phases were produced by repeated sampling of 20 independent data splits. Table C.9 (Appendix C) reports the means and standard deviations of the trials. For visual reference, Figure 3 displays the boxplots of the distributions of the performance indexes of the modalities, grouped by patients’ representation strategy and deconfusion approach. Pairwise tests were performed between settings to be compared and can be appreciated in Table 2.

**Figure 3:**
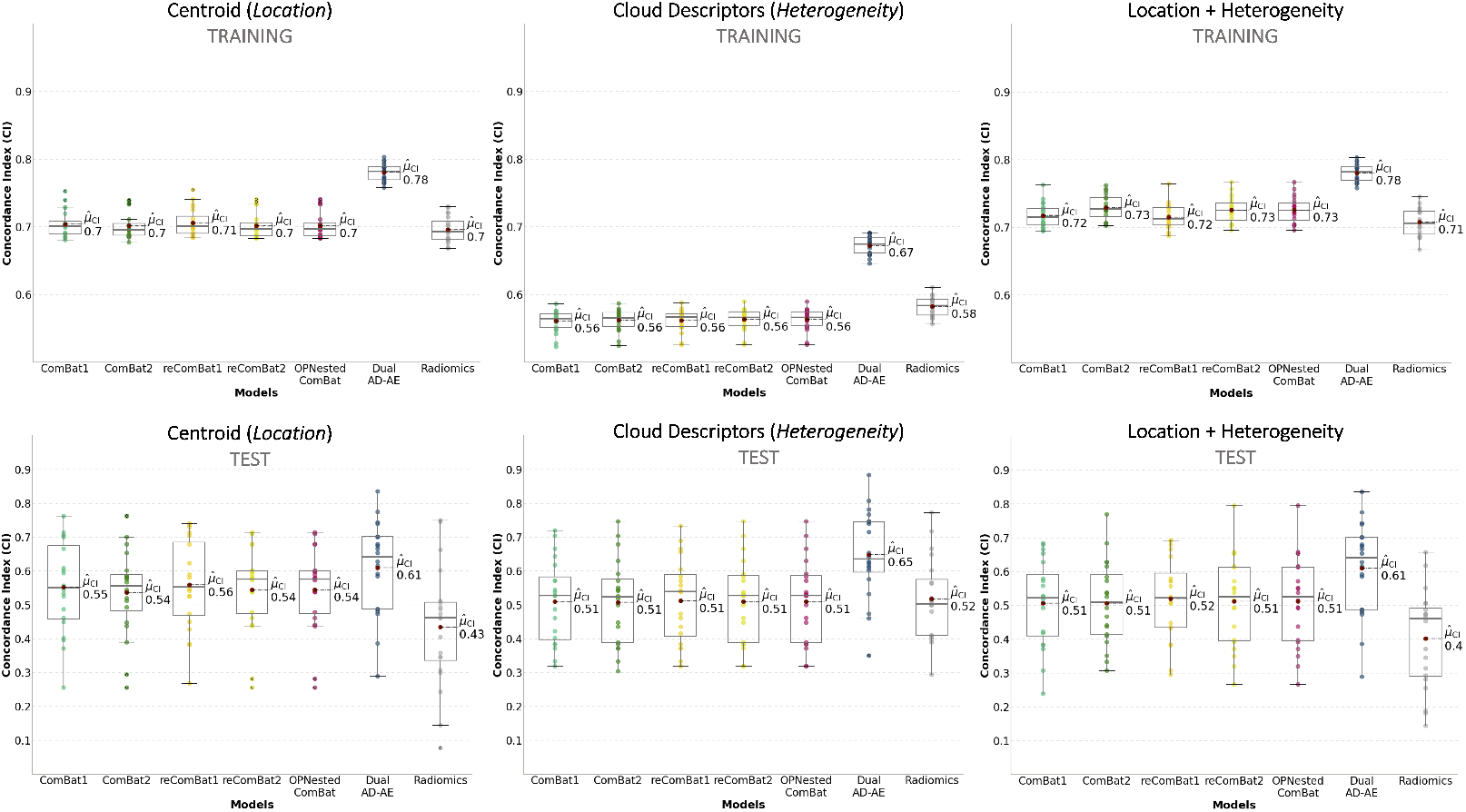
Experiment 3 results: the boxplots of the distributions of algorithms’ performances. The three different patients’ representation strategies are considered per modality and one representation is displayed per plot. The top row plots show training results while the bottom row plots show testing performances. All plots report on the y-axis the CIs of ComBat-center-scanner radiomics (light green, ComBat1 for short), ComBat-scanner-center radiomics (dark green, ComBat2 for short), ReComBat-center-scanner radiomics (dark yellow, reComBat1 for short), ReComBat-scanner-center radiomics (light yellow, reComBat2 for short), and OPNested ComBat radiomics (fuchsia), dual AD-AE embeddings (blue) and original radiomics (grey). The red dots highlight the mean CIs, which are also reported on the right of each respective boxplot.

**Table 2:**
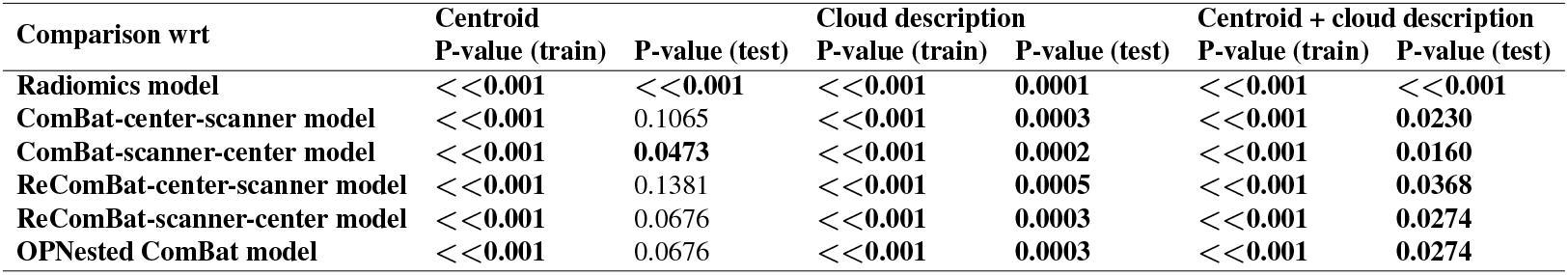
Experiment 3 results: p-values of the tests comparing different survival models. The Dual AD-AE model is considered according to its three different patient representations: (1) the patient is described by the centroid of its point cloud, (2) the patient is described by the topological characteristics of its point cloud and (3) the patient is described by both the centroid and the topological characteristics of its point cloud. These are compared with the radiomics-based models, the ComBat-based models, the ReComBat-based models, and with the OPNested-based models. Comparisons are made upon the same patient representation: for instance, the Dual AD-AE model fed with centroid representation is compared to the models described in the columns which were fed with centroid representation as well, and so on. Significant values are highlighted in bold.

As displayed in Table 2, the model performance of the Dual AD-AE modality was significantly higher than radiomics’, suggesting how the deconfusion step does also benefit the prediction and the signal-to-noise ratio. Of note, the patient representations including cloud topology descriptors (i.e. when using heterogeneity as a predictor) always achieved better performance than the benchmarks, being the most predictive and generalizable (i.e. test set performance) overall. For what centroid representation is concerned, ComBat-center-scanner, ReComBat-center-scanner, ReComBat-scanner-center, and OPNested ComBat scored similarly to the Dual AD-AE. We remind that the OPNested algorithm implemented the same sequence of ComBat-scanner-center, however, this latter model had lower performance, being significantly outperformed by our model.

### 3.4. Alternative to deconfusion: frailty Cox proportional hazards model

Deconfusion methods ultimately allow the effective modeling of patients’ representations across scanners and centers. However, instead of removing the confounding factor, an alternative yet the well-established approach to model multi-source samples (i.e. multi-center/multi-scanner data, where we have dependence within groups) is the explicit modeling of the group-specific variability within the prediction model. For time-to-event data, this can be done via the frailty Cox proportional hazards model (Kosorok et al., 2004), which estimates center and/or scanner random effects together with the baseline hazard function. To verify whether this approach would make the deconfusion step irrelevant, the centroid representation derived from the raw radiomics features was reduced by PCA and fed into a frailty Cox proportional hazard model with center-specific and scanner-specific random intercepts. Unfortunately, this test dramatically failed due to a lack of model convergence. This result is motivated by the small sample size of the data at hand, combined with the high dimensionality of the radiomic variable (even after PCA) and the large number of censored patients, which did not allow the model to properly estimate the effects’ parameters neither on the training sets, on the testing sets, nor the dataset as a whole.

## 4. Discussion

In this work, we developed a deconfusion algorithm to harmonize multi-center imaging data, with a particular focus on multi-lesion/metastatic cancers, such as Hodgkin Lymphoma. The Dual AD-AE model performed dimensionality reduction of radiomic features while removing center- and scanner-related information simultaneously. The proposed approach was trained on a dataset of Hodgkin Lymphoma patients from two centers and outperformed the state-of-the-art methods in the prediction of response to first-line chemotherapy.

Three experiments were performed to evaluate the model’s properties, raising some major points of discussion. First, the deconfounding power of the Dual AD-AE was granted. In fact, the accuracy of Logistic Regression models predicting the scanner and the center target variable sensibly decreased after deconfusion. The Dual AD-AE demonstrated a comparable deconfusion power to ComBat-based models, showing no statistical differences in cross-validation. However, removing both confounding factors at the same time may uncover and discard inter-confounder relationships which may contribute to undesirable noise in the signal. Interestingly, the standard deviation of the accuracy of the dual AD-AE model in predicting the scanner type was lower than other models, suggesting the robustness and stability of the proposed model. The ComBat (and ReComBat) algorithm applied twice showed variable results when changing the order of application. This inconsistency is not surprising, as it motivated the development of OPNested ComBat in the first place (Horng et al., 2022a,b). Despite the slight algorithmic differences between ComBat and OPNested, OPNested performed very similarly to ComBat-scanner-center.

Additionally, as the context of multi-lesion/metastatic data may benefit from the exploitation of intra-tumor heterogeneity as predictive information, we designed a novel metric (PCSD) and an associated empirical test to quantify the impact of AD-AEs deconfusion and dimensionality reduction on intra-lesion relationships shaping the spatial conformation of patients’ point clouds. Overall, the Dual AD-AE resulted in a significantly low PCSD value, rejecting the null hypothesis of no correlation between the original (raw radiomics) and the deconfounded clouds of lesions. On one hand, this was expected and desired as lesions of one patient share both the same center and scanner variability, that is, noise can be considered constant within a single patient. Indeed, the relationship among peer lesions should in principle not be spoiled by center and scanner deconfusion. On the other hand, it might be possible that minor shifts could be appreciated in specific lesions, especially where massive non-linear transformations were needed to properly clear the data. This might be true for some patients lying on the far-right tail of the PCSD population distribution. As proved by the test, such results do not translate into a detrimental data transformation, rather they show that a trade-off between deconfusion and cloud-shape invariance has to be tuned and rigorously assessed. On purpose, the PCSD metric can be exploited to highlight the presence of such additional sources of latent and interactive noise, that once removed would release the true predictive power of intra-lesion heterogeneity.

This point was further validated in the third experiment presented in this work, where we assessed the increase in the prognostic power of the deconfounded representation of patients in terms of response to therapy, against ComBat-based alternative approaches. In principle, a proper deconfusion allows the shape and location of the point clouds coming from different sources to be meaningfully compared. Thus, one can expect that predictive models built on these clouds’ representation, that is lesions’ characteristics and intra-tumor heterogeneity, benefit from the deconfusion process. Dual AD-AE embeddings showed significant improvements compared to the baseline and the benchmarks. This testifies how the proposed model can identify and remove the complex and potentially nonlinear portion of confounders’ noise that the competitors ignore. Moreover, it demonstrates the relevance of removing all confounders simultaneously when in presence of multiple factors of variance in the data.

A further particularly relevant result was the difference in performance when using heterogeneity (i.e. cloud description indexes) as a predictor. While this cloud shape representation was merely a simple proof-of-concept example, Dual AD-AE embedding was seen to allow for a much better prediction than the baseline model and competitors. Conversely, ComBat-based and ReComBat-based benchmarks seemed to corrupt the heterogeneity signal to the point of achieving lower CI than the original radiomics features during training, and they grant just a very limited performance increase during testing. Additionally, to the best of our knowledge, none of the previous studies comparing deconfusion algorithms for radiomics data (Da-Ano et al., 2020, Ligero et al., 2021, Mali et al., 2021) evaluated their impact on the predictive power of groups of lesions. Here, our proposed approach was the only deconfounding algorithm truly releasing the predictive power of heterogeneity, which became the most generalizable predictor.

This finding leads to two relevant considerations. Clinically speaking, it supports the hypothesis that intra-lesion heterogeneity does carry predictive information, once properly corrected for linear and non-linear confounders. Technically, it endorses the use of a more complex non-linear model like the AD-AE, that can uncover and remove explicit and latent types of noise effectively. Although not explicitly enforcing inter-lesion relationships consistency in the model we propose, so that it could be in principle applied as-is to single-lesion data, this result testifies in favor of its application (as opposed to the state-of-the-art) to contexts in which heterogeneity information is crucial for prediction.

Of course, training complex, non-linear, and heavily parametrized models such as the Dual AD-AE has higher computational, time, and memory demands compared to the simpler ComBat-based methods. Nevertheless, the latter algorithms rely on Gaussian distribution assumptions for estimating the parametric definitions of the statistical moments across batches (i.e. the mean and the variance across centers or scanners), prior to standardization. However, this strong hypothesis of underlying data structure may not always be appropriate for radiomics data, leading to underpowered and biased transformations. Conversely, we proposed a non-parametric algorithm removing linear and non-linear confounder-induced noise without any prior assumption. Furthermore, the Dual AD-AE was the only method that dealt with two confounders simultaneously. This permitted the reduction of the risk of ignoring the portion of noise induced by center and scanner interactions (for instance, if one center uses way more frequently a set of parameters for a specific scanner, compared to other centers). Moreover, thanks to its modular nature, one could easily extend the model to adversarially predict - that is, unlearn - more than two confounders. Additional branches could be added, and the overall loss in Equation 1 might be updated with the maximization of the corresponding accuracies. Further, the weighting parameters *λ_i_* (with *i* being the number of adversary branches) enable defining the impact of each confounder, rebalancing the expected (or measured) relative effect of noising factors on the data. Both these aspects could hardly be integrated into the ComBat approach. Finally, as opposed to ComBat-based methods, Dual AD-AE performs dimensionality reduction together with cleaning the embeddings. While this may affect the interpretability of the deconfounded data, we argue that radiomics features are not easily interpretable *per se*, and they usually need a dimensionality reduction (such as PCA) before modeling, as they are highly collinear.

Additionally, we tested the performance of multi-level models as an alternative to deconfusion. Indeed, disregarding the deconfounding algorithm employed, the two-step pipeline of removing confounding effects and then analyzing the corrected data has raised several critiques (Nygaard et al., 2016, Zindler et al., 2020). Oppositely, the most sponsored solution when the confounder information is available is including it within the final prediction model. Nevertheless, we have shown in our last tentative experiment how a frailty CoxPH model (even if with only one confounder) does not converge when the sample size is small and the number of censored patients is high. This is quite common in multi-center studies of rare diseases.

A limitation of the present study is the lack of further data to test our proposed approach. However, no additional comparable data was available to the authors at the time of writing. Nevertheless, we believe that the comprehensive tests and benchmark studies performed on these cohorts represent a valuable proof-of-concept of the method’s potential.

In conclusion, we provided a modular and effective approach for harmonizing imaging data coming from different sources. We proved that our approach could efficiently correct for multiple batch-related differences so that data appear as if they were acquired under a common set of conditions. This translates to higher prognostic performances, above all for what regards intra-tumor heterogeneity of multi-lesions/metastatic cancers. Finally, it is well known that NN models such as the Dual AD-AE can benefit from Transfer Learning (Tan et al., 2018) to aid the problem of suboptimal and/or overfitting parameters when training data is limited. Therefore, we provide a tutorial to apply our method to new data, available in We currently share the weights of our pre-trained network on this study’s [GitHub](https://github.com/LaraCavinato/Dual-ADAE). We currently share the weights of our pre-trained network on this study’s cohorts. Researchers might thus decide to use such weights to pre-train their Dual AD-AE model, “borrowing” information from additional samples without privacy concerns. This model-sharing framework could be pushed forward with the contribution of the scientific community sharing their fine-tuned parameters, paving the way for a virtuous cycle of open science. Insightful knowledge could be thus derived from more exhaustive models to optimally impact the clinical practice.

## 5. Conclusions

In this work, we developed a Dual AD-AE algorithm for harmonizing multi-center imaging data, with a focus on multi-lesion/metastatic cancers like Hodgkin Lymphoma. The proposed model performed dimensionality reduction of radiomic features while removing center- and scanner-related information. The algorithm was trained on a dataset of Hodgkin Lymphoma patients from two centers and out-performed state-of-the-art methods in predicting response to first-line chemotherapy. Three experiments were performed to evaluate the algorithm’s properties, which showed its deconfounding power and robustness compared to ComBat-based models. We also introduced a novel metric (PCSD) to quantify the impact of AD-AEs on intra-lesion relationships. The Dual AD-AE resulted in a low PCSD value and improved the prognostic power of the deconfounded representation of patients compared to ComBat-based models. The Dual AD-AE also showed better prediction performance using heterogeneity as a predictor. This supports the hypothesis that intra-lesion heterogeneity carries predictive information and endorses the use of a more comprehensive deconfounding algorithm for radiomics data.

## Appendix A. Patients’ characteristics

**Table A.3:**
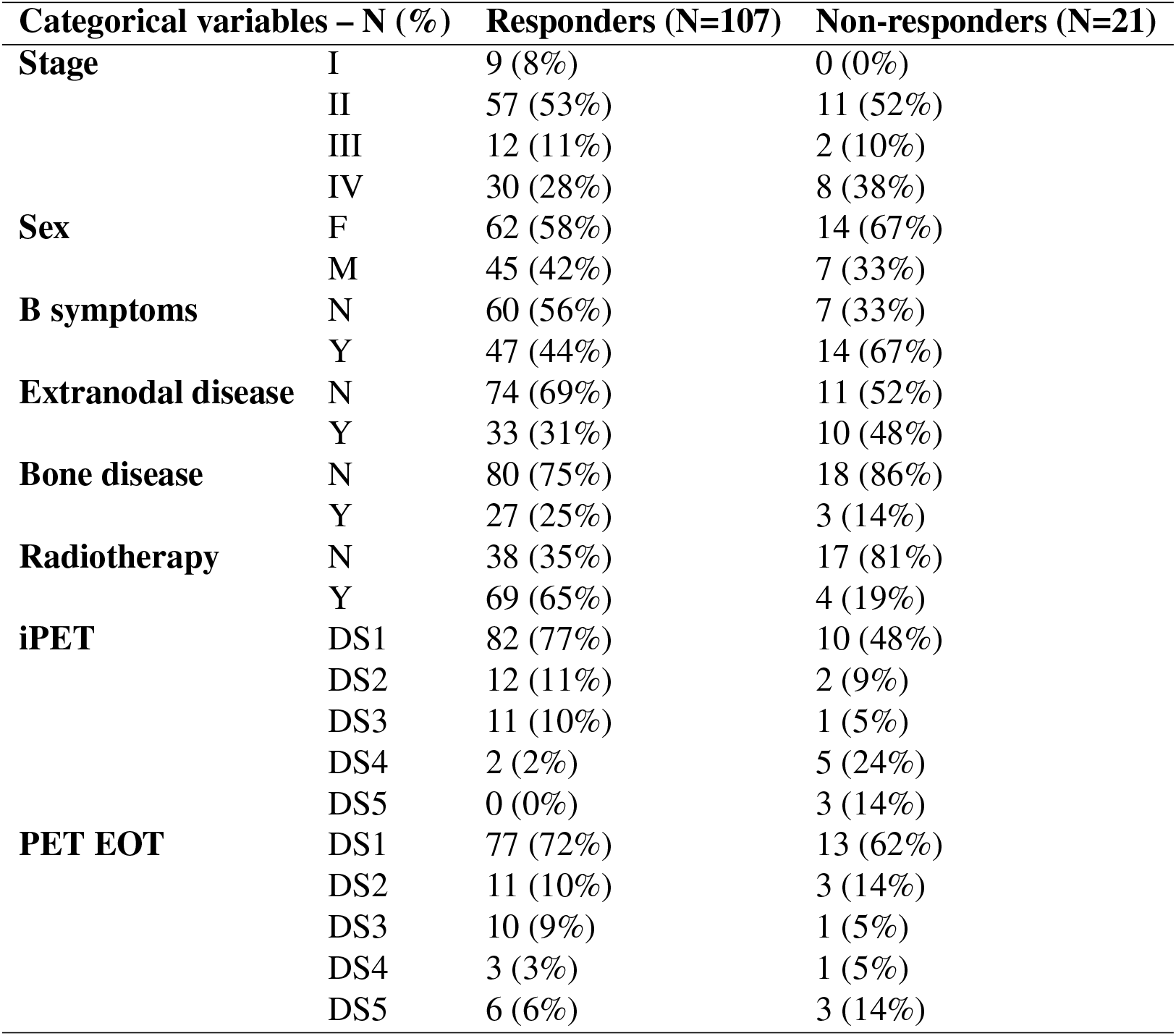
Patients’ characteristics in Institution 1: variables are divided into categorical (number, percentage on the total) and numerical (mean, standard deviation). In the first group, they are listed the stage (four statuses), the sex (female F and male M), the presence of B symptoms like fever, sweats, weight loss (yes Y and no N), status of the disease (extranodal disease: yes Y and no N; bone disease: yes Y and no N), administration of radiotherapy (yes Y and no N), the outcome of interim PET (iPET, Deauville Score DS of the PET), end of treatment PET (EOT PET, Douville Score DS of the PET). Statistics are stratified by the treatment response, thus patients are divided into responders and non-responders.

**Table A.4:** Patients’ characteristics in Institution 1: variables are divided into categorical (number, percentage on the total) and numerical (mean, standard deviation). Among the numerical variables, there are age, number of nodal lesions of the patients, number of extranodal lesions of the patients, and time to relapse (for censored patients, the time to last follow-up is taken). Statistics are stratified by the treatment response, thus patients are divided into responders and non-responders.

| **Numerical variables – mean (std deviation)** |  |  |
| --- | --- | --- |
| Age | 39.252 (15.875) | 40.143 (15.963) |
| # Nodal lesions | 6.673 (4.813) | 6.619 (6.184) |
| # Extranodal lesions | 1.916 (5.750) | 3.857 (10.256) |
| Time to relapse [days] | 1126.97 (704.94) | 358.86 (322.854) |

**Table A.5:** Patients’ characteristics in Institution 2: variables are divided into categorical (number, percentage on the total) and numerical (mean, standard deviation). In the first group, they are listed the stage (four statuses), the sex (female F and male M), the presence of B symptoms like fever, sweats, weight loss (yes Y and no N), status of the disease (extranodal disease: yes Y and no N; bone disease: yes Y and no N), administration of radiotherapy (yes Y and no N), the outcome of interim PET (iPET, positive or negative), end of treatment PET (EOT PET, positive or negative). Statistics are stratified by the treatment response, thus patients are divided into responders and non-responders.

| **Categorical variables – N (%)** |  | **Responders (N=59)** | **Non-responders (N=17)** |
| --- | --- | --- | --- |
| **Stage** | I | 1 (2%) | 0 (0%) |
|  | II | 31 (52%) | 4 (23%) |
|  | III | 6 (10%) | 1 (6%) |
|  | IV | 21 (36%) | 12 (71%) |
| **Sex** | F | 34 (58%) | 8 (47%) |
|  | M | 25 (42%) | 9 (53%) |
| **B symptoms** | N | 35 (59%) | 4 (23%) |
|  | Y | 24 (41%) | 13 (77%) |
| **Extranodal disease** | N | 39 (65%) | 7 (41%) |
|  | Y | 20 (45%) | 10 (59%) |
| **Bone disease** | N | 44 (75%) | 13 (77%) |
|  | Y | 15 (25%) | 4 (23%) |
| **Radiotherapy** | N | 20 (45%) | 14 (82%) |
|  | Y | 39 (65%) | 3 (18%) |
| **iPET** | Negative | 55 (93%) | 8 (47%) |
|  | Positive | 4 (7%) | 9 (53%) |
| **PET EOT** | Negative | 59 (100%) | 0 (0%) |
|  | Positive | 0 (0%) | 17 (100%) |

**Table A.6:** Patients’ characteristics in Institution 2: variables are divided into categorical (number, percentage on the total) and numerical (mean, standard deviation). Among the numerical variables, there are age, number of nodal lesions of the patients, number of extranodal lesions of the patients, and time to relapse (for censored patients, the time to last follow-up is taken). Statistics are stratified by the treatment response, thus patients are divided into responders and non-responders.

| **Numerical variables – mean (std deviation)** |  |  |
| --- | --- | --- |
| Age | 36.478 (13.915) | 42.867 (17.868) |
| # Nodal lesions | 7.271 (5.499) | 9.706 (6.362) |
| # Extranodal lesions | 2.288 (5.789) | 3.706 (7.355) |
| Time to relapse [days] | 1105.72 (546.490) | 257.59 (167.17) |

## Appendix B. Scanners’ specifications

**Table B.7:** Image acquisition protocols and scanner specification in Institution 1: 85 patients were scanned with Siemens Biograph scanner; 51 patients were scanned with General Electric Discovery 690 scanner; 5 were scanned with other unspecified scanners.

| Institution 1 | Biograph – Siemens |  | Discovery 690 – General Electric |  |
| --- | --- | --- | --- | --- |
|  | PET | CT | PET | CT |
| **Min/bed position (static/dynamic)** | 2.5 (static) | – | 2 (static) | – |
| **Crystal** | LSO | – | LYSO | – |
| **Reconstruction** | Iterative | – | Iterative, TOF Sharp IR | – |
| **Attenuation correction** | On CT data | – | On CT data | – |
| **Matrix (pixels)** | 128×128 | 512×512 | 256×256 | 512×512 |
| **Resolution (mm)** | 5.3×5.3 | 0.98×0.98 | 2.73×2.73 | 1.37×1.37 |
| **Slice thickness (mm)** | 2.0 | 4.0 | 3.27 | 3.27 |
| **Slices** | – | 6 | – | 64 |

**Table B.8:**
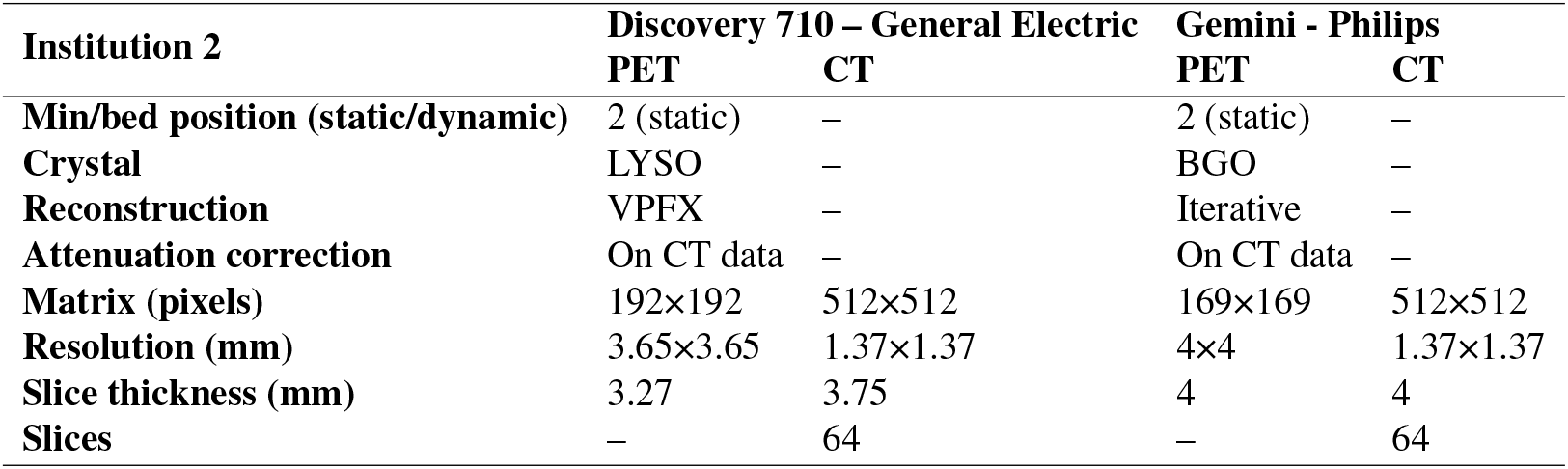
Image acquisition protocols and scanner specification in Institution 2: 34 patients were scanned with General Electric Discovery 710 scanner; 38 patients were scanned with Philips Gemini scanner; 1 patient was scanned with other unspecified scanners.

| Institution 2 | Discovery 710 – General Electric |  | Gemini - Philips |  |
| --- | --- | --- | --- | --- |
|  | PET | CT | PET | CT |
| **Min/bed position (static/dynamic)** | 2 (static) | – | 2 (static) | – |
| **Crystal** | LYSO | – | BGO | – |
| **Reconstruction** | VPFX | – | Iterative | – |
| **Attenuation correction** | On CT data | – | On CT data | – |
| **Matrix (pixels)** | 192×192 | 512×512 | 169×169 | 512×512 |
| **Resolution (mm)** | 3.65×3.65 | 1.37×1.37 | 4×4 | 1.37×1.37 |
| **Slice thickness (mm)** | 3.27 | 3.75 | 4 | 4 |
| **Slices** | – | 64 | – | 64 |

## Appendix C. Results of experiment 3

**Table C.9:** Experiment 3 results: performance of the Cox proportional hazard models trained and tested in cross-validation using different patient representations. Each modality – i.e. radiomics, Dual AD-AE embeddings, ComBat-based standardization, ReComBat-based standardization, and OPNested-based standardization – is fed in the survival model according to three different patient representations: (1) the patient is described by the centroid of its point cloud (“centroid representation”), (2) the patient is described by the topological characteristics of its point cloud (“cloud description representation”) and (3) the patient is described by both the centroid and the topological characteristics of its point cloud (“complete representation”). Best values are highlighted in bold.

| Models | Centroid representation |  | Cloud description representation |  | Complete representation |  |
| --- | --- | --- | --- | --- | --- | --- |
|  | C-index (train) | C-index (test) | C-index (train) | C-index (test) | C-index (train) | C-index (test) |
| Radiomics | 0.6962 ± 0.0190 | 0.4344 ± 0.1779 | 0.5828 ± 0.0141 | 0.5184 ± 0.1203 | 0.7077 ± 0.0207 | 0.4026 ± 0.1460 |
| Dual AD-AE | **0.7803 ± 0.0132** | **0.6100 ± 0.1382** | **0.6728 ± 0.0135** | **0.6481 ± 0.1305** | **0.7803 ± 0.0132** | **0.6100 ± 0.1382** |
| ComBat-center-scanner | 0.7041 ± 0.0201 | 0.5525 ± 0.1358 | 0.5611 ± 0.0162 | 0.5101 ± 0.1228 | 0.7173 ± 0.0180 | 0.5063 ± 0.1279 |
| ComBat-scanner-center | 0.7016 ± 0.0195 | 0.5366 ± 0.1266 | 0.5625 ± 0.0159 | 0.5075 ± 0.1262 | 0.7291 ± 0.0185 | 0.5066 ± 0.1234 |
| ReComBat-center-scanner | 0.7064 ± 0.021 | 0.5595 ± 0.1344 | 0.5621 ± 0.0161 | 0.5128 ± 0.1239 | 0.7153 ± 0.0186 | 0.5201 ± 0.1224 |
| ReComBat-scanner-center | 0.7019 ± 0.0194 | 0.5444 ± 0.1240 | 0.5635 ± 0.0160 | 0.5101 ± 0.1244 | 0.7260 ± 0.0192 | 0.5120 ± 0.1348 |
| OPNested ComBat | 0.7019 ± 0.0194 | 0.5444 ± 0.1240 | 0.5635 ± 0.0160 | 0.5101 ± 0.1244 | 0.7260 ± 0.0192 | 0.5120 ± 0.1348 |

## Acknowledgment

We acknowledge all the personnel of the Nuclear Medicine Department of IRCCS Humanitas Clinical and Research Hospital and Fondazione IRCCS Istituto Nazionale dei Tumori for the assistance during the PET/CT scans, segmentation of lesions, extraction of radiomic features and retrieval of patients’ personal information from electronic health records.

## References

Adamer, M.F., Brüningk, S.C., Tejada-Arranz, A., Estermann, F., Basler, M., Borgwardt, K., 2022. recombat: batch-effect removal in large-scale multi-source gene-expression data integration. Bioinformatics Advances 2, vbac071.

Afshar, P., Mohammadi, A., Plataniotis, K.N., Oikonomou, A., Benali, H., 2019. From handcrafted to deep-learning-based cancer radiomics: challenges and opportunities. IEEE Signal Processing Magazine 36, 132–160.

Berenguer, R., Pastor-Juan, M.D.R., Canales-Vázquez, J., Castro-García, M., Villas, M.V., Mansilla Legorburo, F., Sabater, S., 2018. Radiomics of ct features may be nonreproducible and redundant: influence of ct acquisition parameters. Radiology 288, 407–415.

Campbell, P.J., Pleasance, E.D., Stephens, P.J., Dicks, E., Rance, R., Goodhead, I., Follows, G.A., Green, A.R., Futreal, P.A., Stratton, M.R., 2008. Subclonal phylogenetic structures in cancer revealed by ultra-deep sequencing. Proceedings of the National Academy of Sciences 105, 13081–13086.

Cavinato, L., Pegoraro, M., Ragni, A., Ieva, F., 2022. Imaging-based representation and stratification of intra-tumor heterogeneity via tree-edit distance. Scientific reports 12, 19607.

Chen, C., Grennan, K., Badner, J., Zhang, D., Gershon, E., Jin, L., Liu, C., 2011. Removing batch effects in analysis of expression microarray data: an evaluation of six batch adjustment methods. PloS one 6, e17238.

Da-Ano, R., Visvikis, D., Hatt, M., 2020. Harmonization strategies for multicenter radiomics investigations. Physics in Medicine & Biology 65, 24TR02.

Dincer, A.B., Janizek, J.D., Lee, S.I., 2020. Adversarial deconfounding autoen-coder for learning robust gene expression embeddings. Bioinformatics 36, i573–i582.

Gil, D., Ramos, O., Perez, R., 2021. Topological radiomics (topiomics): Early detection of genetic abnormalities in cancer treatment evolution, in: Extended Abstracts GEOMVAP 2019: Geometry, Topology, Algebra, and Applications; Women in Geometry and Topology, Springer. pp. 89–93.

Harrell, F.E., Califf, R.M., Pryor, D.B., Lee, K.L., Rosati, R.A., 1982. Evaluating the yield of medical tests. Jama 247, 2543–2546.

Horng, H., Singh, A., Yousefi, B., Cohen, E.A., Haghighi, B., Katz, S., Noël, P.B., Kontos, D., Shinohara, R.T., 2022a. Improved generalized combat methods for harmonization of radiomic features. Scientific Reports 12, 19009.

Horng, H., Singh, A., Yousefi, B., Cohen, E.A., Haghighi, B., Katz, S., Noël, P.B., Shinohara, R.T., Kontos, D., 2022b. Generalized combat harmonization methods for radiomic features with multi-modal distributions and multiple batch effects. Scientific reports 12, 1–12.

Johnson, W.E., Li, C., Rabinovic, A., 2007. Adjusting batch effects in microarray expression data using empirical bayes methods. Biostatistics 8, 118–127.

Kosorok, M.R., Lee, B.L., Fine, J.P., 2004. Robust inference for univariate proportional hazards frailty regression models.

Lavin, R.C., Tan, S., 2022. Spatial relationships of intra-lesion heterogeneity in mycobacterium tuberculosis microenvironment, replication status, and drug efficacy. PLoS Pathogens 18, e1010459.

Ligero, M., Jordi-Ollero, O., Bernatowicz, K., Garcia-Ruiz, A., Delgado-Muñoz, E., Leiva, D., Mast, R., Suarez, C., Sala-Llonch, R., Calvo, N., et al., 2021. Minimizing acquisition-related radiomics variability by image resampling and batch effect correction to allow for large-scale data analysis. European radiology 31, 1460–1470.

Lin, D.Y., Wei, L.J., 1989. The robust inference for the cox proportional hazards model. Journal of the American statistical Association 84, 1074–1078.

Mali, S.A., Ibrahim, A., Woodruff, H.C., Andrearczyk, V., Müller, H., Primakov, S., Salahuddin, Z., Chatterjee, A., Lambin, P., 2021. Making radiomics more reproducible across scanner and imaging protocol variations: a review of harmonization methods. Journal of personalized medicine 11, 842.

Mohty, R., Dulery, R., Bazarbachi, A.H., Savani, M., Hamed, R.A., Bazarbachi, A., Mohty, M., 2021. Latest advances in the management of classical hodgkin lymphoma: The era of novel therapies. Blood Cancer Journal 11, 126.

Nioche, C., Orlhac, F., Boughdad, S., Reuzé, S., Goya-Outi, J., Robert, C., Pellot-Barakat, C., Soussan, M., Frouin, F., Buvat, I., 2018. Lifex: a freeware for radiomic feature calculation in multimodality imaging to accelerate advances in the characterization of tumor heterogeneity. Cancer research 78, 4786–4789.

Nygaard, V., Rødland, E.A., Hovig, E., 2016. Methods that remove batch effects while retaining group differences may lead to exaggerated confidence in downstream analyses. Biostatistics 17, 29–39.

Parmar, C., Barry, J.D., Hosny, A., Quackenbush, J., Aerts, H.J., 2018. Data analysis strategies in medical imagingdata science designs in medical imaging. Clinical cancer research 24, 3492–3499.

Pavic, M., Bogowicz, M., Würms, X., Glatz, S., Finazzi, T., Riesterer, O., Roesch, J., Rudofsky, L., Friess, M., Veit-Haibach, P., et al., 2018. Influence of inter-observer delineation variability on radiomics stability in different tumor sites. Acta Oncologica 57, 1070–1074.

Rizzo, A., Triumbari, E.K.A., Gatta, R., Boldrini, L., Racca, M., Mayerhoefer, M., Annunziata, S., 2021. The role of 18 f-fdg pet/ct radiomics in lymphoma. Clinical and Translational Imaging, 1–10.

Sangaletti, S., Iannelli, F., Zanardi, F., Cancila, V., Portararo, P., Botti, L., Vacca, D., Chiodoni, C., Di Napoli, A., Valenti, C., et al., 2020. Intra-tumour heterogeneity of diffuse large b-cell lymphoma involves the induction of diversified stroma-tumour interfaces. EBioMedicine 61, 103055.

Scapicchio, C., Gabelloni, M., Barucci, A., Cioni, D., Saba, L., Neri, E., 2021. A deep look into radiomics. La radiologia medica 126, 1296–1311.

Sollini, M., Kirienko, M., Cavinato, L., Ricci, F., Biroli, M., Ieva, F., Calderoni, L., Tabacchi, E., Nanni, C., Zinzani, P.L., et al., 2020. Methodological framework for radiomics applications in hodgkin’s lymphoma. European journal of hybrid imaging 4, 1–17.

Tabanelli, V., Melle, F., Motta, G., Mazzara, S., Fabbri, M., Corsini, C., Gerbino, E., Calleri, A., Sapienza, M.R., Abbene, I., et al., 2020. Evolutionary cross-roads: morphological heterogeneity reflects divergent intra-clonal evolution in a case of high-grade b-cell lymphoma. Haematologica 105, e432.

Tan, C., Sun, F., Kong, T., Zhang, W., Yang, C., Liu, C., 2018. A survey on deep transfer learning, in: Artificial Neural Networks and Machine Learning–ICANN 2018: 27th International Conference on Artificial Neural Networks, Rhodes, Greece, October 4-7, 2018, Proceedings, Part III 27, Springer. pp. 270–279.

Yao, Y., Rosasco, L., Caponnetto, A., 2007. On early stopping in gradient descent learning. Constructive Approximation 26, 289–315.

Zindler, T., Frieling, H., Neyazi, A., Bleich, S., Friedel, E., 2020. Simulating combat: how batch correction can lead to the systematic introduction of false positive results in dna methylation microarray studies. BMC bioinformatics 21, 1–15.

